# Microclimatic effects on alpine plant and flower visitor communities and their interactions

**DOI:** 10.1101/646752

**Authors:** Lisa-Maria Ohler, Martin Lechleitner, Robert R. Junker

## Abstract

High-alpine ecosystems are commonly assumed to be particularly endangered by climate warming. Recent research, however, suggests that the heterogeneous topography of alpine landscapes provide microclimatic niches for alpine plants, which may buffer negative effects. Whether the microclimatic heterogeneity also affects higher trophic levels remains unknown. This study shows that the variation in mean seasonal soil temperature within a single alpine pasture is within the same range as in plots differing in nearly 500 m in elevation. This pronounced heterogeneity affected the spatial distribution of plant cover, richness of flowering plant species and plant species composition. These microclimatic effects on plant communities also affected richness of flower visiting insects and the frequency and structure of plant-insect interactions suggesting an effect of microclimate also on higher trophic levels. Our results may stimulate a re-evaluation of the consequences of climate warming on ecosystems that may compensate warming by microclimatic refuges.

## Introduction

Alpine ecosystems are particularly sensitive to climate change as scenarios predict severe warming for high elevations in alpine regions (Pepin et al., 2015), temperature-limited alpine plant species are thus threatened by increased temperatures (Dullinger et al., 2012; Jurasinski & Kreyling, 2007). Recently, considerable work has described and tested possible scenarios for responses of alpine plants to climate change (Alexander et al., 2018; Graae et al., 2018). Regional plant diversity in alpine habitats is expected to change as a result of climate warming due to shifts in climate niches and species’ distributions, which may result in increased competition due to shifts of plants’ distributions to higher elevations and consequently the assembly of new communities (Löffler & Pape, 2008; Pauli et al., 2012; Spasojevic, Bowman, Humphries, Seastedt, & Suding, 2013). Plant species richness and composition affects organisms and processes across trophic levels (Scherber et al., 2010). For instance, the diversities of plants and their flower visitors have been shown to be particularly related due to insect-specific preferences for plant species, which is an prerequisite for efficient pollination and thus plant reproduction (Ebeling, Klein, Schumacher, Weisser, & Tscharntke, 2008; Junker et al., 2013; Junker, Blüthgen, & Keller, 2015). Therefore, a change in plant abundance and distribution and shifts in phenologies due to rising temperatures potentially have negative effects on other trophic levels, in particular on flower visiting insects (Hegland, Nielsen, Lázaro, Bjerknes, & Totland, 2009; Hoiss, Krauss, & Steffan□Dewenter, 2015) and thus impact community structure and put ecosystem functions at risk (Dunne, Harte, & Taylor, 2003).

Among the main drivers of (alpine) plant diversity are climatic conditions, topography, elevation, and biotic interactions (Cavieres et al., 2014; Mayor et al., 2017; Opedal, Armbruster, & Graae, 2015). However, plant community composition and diversity is not only shaped by average environmental and climatic conditions but also by local micro-abiotic filtering (Conti, de Bello, Lepš, Acosta, & Carboni, 2017). Specifically soil temperatures, which in contrast to air temperatures are strongly shaped by the local topography and intake of solar radiation (Lembrechts et al., 2018; Scherrer & Körner, 2010), are thought to affect photosynthetic capacity and growth rates of plants (Arft et al., 1999; Körner, 2003; Wundram, Pape, & Löffler, 2010). Consequently, fine-scale environmental heterogeneity can also shape functional traits as well as community structure of plants (Blonder et al., 2018). However, the relationship of microclimate heterogeneity, species richness and composition remains understudied (Zellweger, De Frenne, Lenoir, Rocchini, & Coomes, 2019) and whether microclimatic heterogeneity also affects higher trophic levels such as flower visitors remains unknown. We hypothesise bottom-up effects of small-scale soil temperature heterogeneity not only on plant communities but also on flower visiting insects and plant-insect interactions. This effect across trophic levels may have implications for the potential of microclimate as buffer for climate change impacts.

This study is aiming to investigate (1) microclimatic differences in root zone temperature of 1.5 × 1.5 m plots within 1.25 ha on a topographically heterogeneous alpine pasture with neglectable differences in elevation, (2) the relationship between soil temperature and plant species communities, as well as effects on (3) flower visitor community richness and (4) plant-insect interactions.

## Material and methods

### Field site and mean seasonal soil temperature

Field work was conducted in the mountain range of the Hohe Tauern in the Austrian Alps. The study site (12°49’21” E 47°07’24” N) was located on an alpine pasture at an elevation of 2,273 m a.s.l. The study site was confined by mountain ridges in the north and east, which resulted in a shorter daily period of direct solar radiation in the eastern part of the study site than in the western part. The pasture was characterized by a heterogeneous topography forming flat hills, which also introduced variation in the time of solar radiation and also in angle of sunlight. Shortly after snow melt in 2016, we established 30 plots (1.5 × 1.5 m) within an area of 1.25 ha. The location of plots was chosen to represent the variability in aspect and slope of the hills (Supplementary Table 2). The maximum elevational difference between plots was 21.90 m, suggesting that microclimatic heterogeneity results from differences in position, exposition and inclination but not from differences in elevation. The mean seasonal soil temperature of each plot was measured with a temperature logger (DS1921G-F5 Thermochron iButtons, Maxime Integrated Products, Sunnyvale, CA, USA) with a resolution of 0.5 °C. Each plot was equipped with a temperature logger that was wrapped in a plastic bag and buried in the centre of the plot at a soil depth of 3 cm (Scherrer & Körner, 2010). The soil temperature was measured continuously every 30 minutes from June 24^th^ 2016 to September 06^th^ 2016 (Supplementary Table 2), which covers the full vegetation period at this elevation. In order to relate the microclimatic heterogeneity within the study site to temperature differences along the elevational gradient in the same Alpine region, we selected eight comparable pastures between 1227 and 2634 m a.s.l and recorded the mean seasonal soil temperature in the same way as described above at up to six positions per elevation. Additionally, we used an infrared thermal camera (mobileIR E9, InfraTec GmbH, Dresden, Germany) equipped with a wide-angle lens (13 mm / 40.53° × 30.96°) to assess the heterogeneity (i.e. standard deviation) in surface temperature of the whole study area. For this we mounted the infrared thermal camera on top of a higher elevated hill in the east of the study area and took an image of the whole study area at noon at the peak of the vegetation season. To investigate the heterogeneity in surface temperature within each of the single study plots we took infrared thermal pictures of every plot seven to nine times a day following a randomized order to account for different weather conditions and then calculated the variations in soil temperature heterogeneity over the course of the day using a quadratic model. Thermal photographs were analysed using the software IRBIS 3plus (mobileIR E9, InfraTec GmbH, Dresden, Germany).

### Vegetation and flower-visitor interactions

Once per week throughout the vegetation season, we recorded the floral abundance of all entomophilous plant species in anthesis per plot (Supplementary Table 3). Floral abundance was defined as the number of floral units (i.e. individual flowers or inflorescences). Thus, the species richness of flowering plants per plot was defined as the total number of flowering entomophilous species per plot throughout the vegetation period. The information on flower abundance per species per plot was also used to compare the community composition of flowering plants between the different plots. Total plant cover per plot in percent was recorded once in the middle of the growing season (Supplementary Table 2). At each plot, flower-visitor interactions were recorded at four events between 07.07.2016 and 18.08.2016 for a five-minute period (Supplementary Table 4). Each sampling day, plots were visited in a randomized order to avoid temporal and spatial biases. The observation of insect visitation was conducted during clear sky conditions. All insects interacting with flowers in anthesis were captured in plastic jars for subsequent determination on family level, if possible on genus or species level (detailed list of taxa provided in Supplementary Table 5). A statistical analysis on the species level is not meaningful due to the high diversity in arthropod species resulting in only one or few observations per species. From these recordings we received the number insect families as well as the total number of insects interacting with flowers for each plot and additionally calculated the complementary specialization *H*_*2*_*’* of the insect-plant network per plot using the R package *bipartite* (Dormann, Fründ, Blüthgen, & Gruber, 2009).

### Statistical analysis

#### Microclimatic heterogeneity relative to temperature differences along an elevational gradient

– In order to relate the microclimatic heterogeneity with the temperature differences along the elevational gradient, we exploited temperature data from two sources. We used our own temperature logger measurements at eight elevations between 1227 and 2636 m a.s.l. (see above) and additionally, we retrieved mean temperature data (1979-2013) from Chelsa (Karger, Conrad, Böhner, Kawohl, Kreft, Soria-Auza, & Kessler, 2017; Karger, Conrad, Böhner, Kawohl, Kreft, Soria-Auza, Zimmermann, et al., 2017) for the months June, July, August, September, which corresponds to the period of the year when we used the temperature loggers. We estimated linear regression models to determine the relationship between elevation and temperature in our study region and used this model to evaluate which differences in elevation correspond to the temperature variations among the microclimatic plots.

#### Flowering plant species composition

To test whether the plant species composition is affected by the mean seasonal soil temperature on micro-plots, we compared the distance matrices based on the quantitative composition of flowering plant species (floral abundance counts, Bray-Curtis distances) and based on the mean seasonal soil temperature measured on the same plots (Euclidean distances) using a Mantel test. Distance matrices and Mantel test were calculated using the R package *vegan* 2.4-2 (Oksanen et al., 2017). Furthermore, the Bray-Curtis distances reflecting the dissimilarity in qualitative plant species composition were visualized using an NMDS calculated with the R package *MASS* 7.3-51.1 (Venables & Ripley, 2002) and temperatures at these plots were added with a thin plate spine surface for interpolating and smoothing the data using the R package *fields* 9.6 (Nychka, Furrer, Paige, & Sain, 2017).

#### Relationship between microclimatic soil temperature on plant and insect richness and their interactions

We tested the pairwise Pearson’s product-moment correlations between the mean seasonal soil temperature, plant cover, species richness of flowering plants, number of insect families recorded on flowers, number of observed insect interactions with flowers and complementary specialization *H*_*2*_*’* of flower-visitor interaction networks. Using Pearson’s correlation coefficient *r* and we established a correlation network using the R package *igraph* 1.2.2 (Csardi & Nepusz, 2006) restricting the analysis to significant correlation (*p* < 0.05) only. From the correlation network, we extracted the betweenness centrality (*BC*) which is defined by the number of shortest paths going through a node and the degree centrality (*DC*) of the nodes which is the number of its adjacent edges. Additionally, using the same data we performed a structural equation model SEM to support the findings form correlation network analysis. However, because the sample size is not suited to perform SEM, we present these data in Supplementary Information 1, only to support the results presented in the main text.

## Results

### Mean seasonal soil temperature

The mean seasonal soil temperature of our study plots measured with temperature loggers in xx cm belowground ranged between 9.60 °C and 12.40 °C (mean ± SD = 10.88 ± 0.75 °C). The largest difference in mean seasonal soil temperature between two study plots was therefore 2.8 °C (Fig. 1). Microclimatic heterogeneity of surface temperature within plots as recorded by an infrared thermal camera was subjected to strong diurnal variations (quadratic model: *F*_2, 239_ = 39.73, *p* < 0.001, *r*^*2*^ = 0.24, Fig. 2a) and peaked between 1400 and 1500 (UTC+02:00) when temperature was distributed most heterogeneously across the plots indicating that solar radiation is the source of microclimatic heterogeneity in our system.

**Fig 1.**
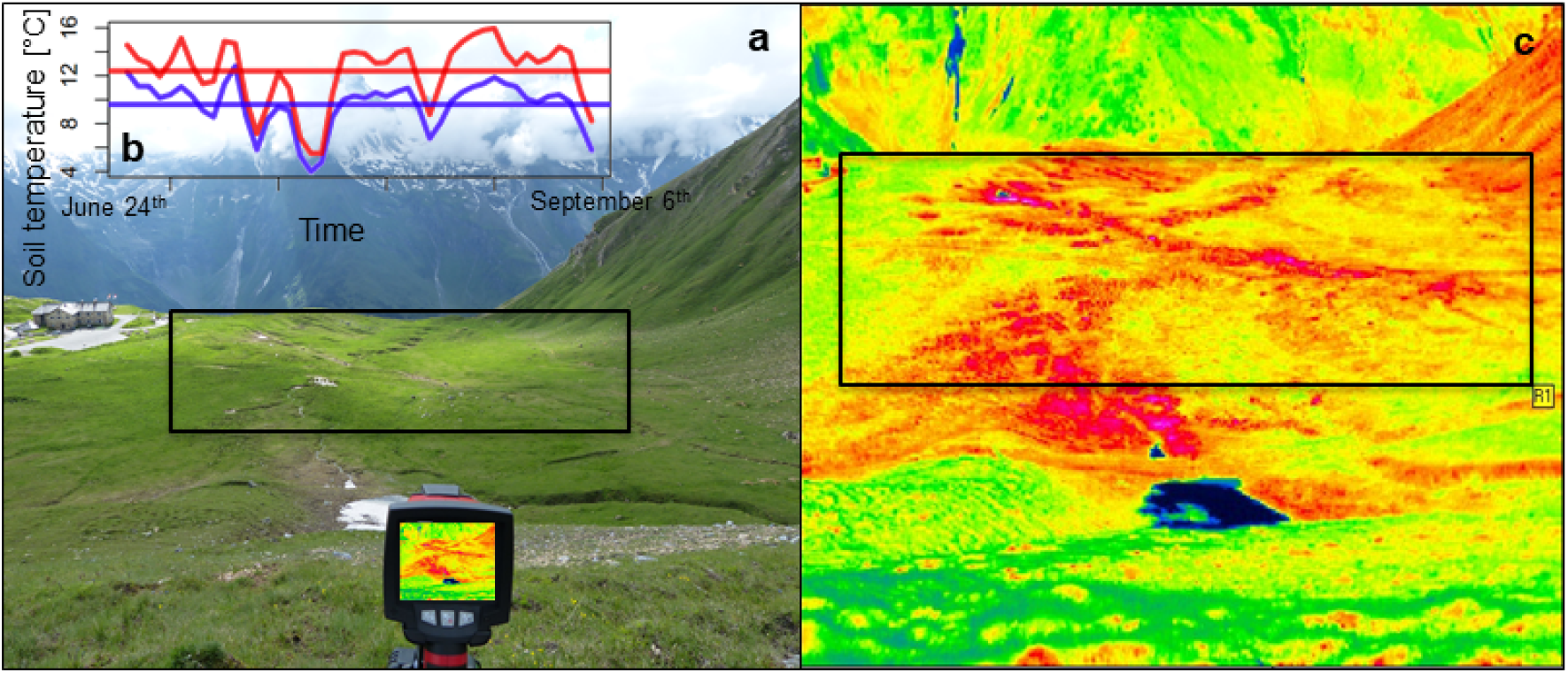
Study site with pronounced microclimatic heterogeneity. **a**, Study site, a high alpine pasture with heterogenous topography located at 2,273 m a.s.l. in the mountain range of the Hohe Tauern in the Austrian Alps (12°49’21” E 47°07’24” N). The *n* = 30 plots were distributed within the black frame. **b**, Seasonal course of soil temperature of the vegetative season in 2016 of the warmest (red) and the coldest plot (blue) measured with temperature loggers. The straight lines indicate the mean seasonal soil temperatures of the warmest (upper line, red) and coldest plot (lower line, blue). **c**, False color image of the study site taken with an infrared thermal camera showing the microclimatic heterogeneity in surface temperature at noon. Each pixel represents the surface temperature of plot a given position. Black to blue pixels represent cold surface temperatures (e.g. the snowfield in the foreground), warmer colours are represented by a gradient from green over yellow to red and pink. The *n* = 30 plots were distributed within the black frame.

**Fig. 2.**
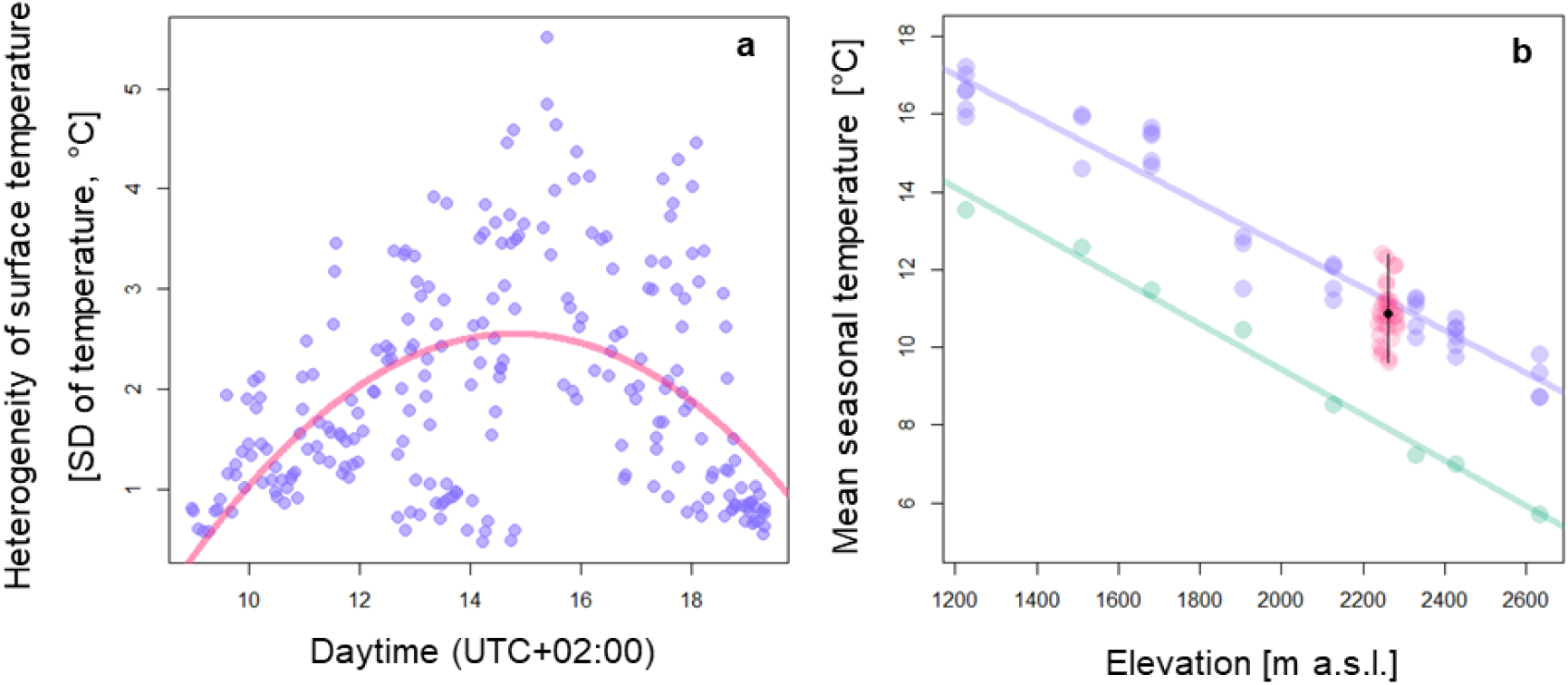
Diurnal course of surface temperature heterogeneity within plots and mean seasonal temperature across plots and spatial scales. **a**, Microclimatic heterogeneity (i.e. standard deviation in surface temperature within plots) of all plots across the study site recorded by an infrared thermal camera following a diurnal course calculated with a quadratic model (red line). *n* = 30 plots (2.25 m^2^) were photographed up to nine times at different times of the day between 0900 and 1930 (UTC+02:00). **b**, Mean seasonal temperature measurements of different sources and scales within the study area. Mean seasonal soil temperature of *n* = 30 plots (red points) on an alpine pasture located at 2,273 m a.s.l. in the mountain range of the Hohe Tauern in the Austrian Alps. Mean seasonal mean soil temperature and standard deviation are given in black. Seasonal mean soil temperature at eight elevations between 1227 and 2634 m a.s.l. in the same area measured with temperature loggers (blue points), regression line is given as blue line. Air temperature data (1979-2013) retrieved from Chelsa climate database for the same eight elevations (green), regression line is given as green line.

Mean seasonal temperature of other pastures between 1227 and 2636 m a.s.l clearly decreased with increasing elevation both based on our own measurements using temperature loggers buried in soil (Pearson’s product-moment correlation: *t*_34_ = −24.84, *p* < 0.001, *r*^*2*^ = 0.95, slope of regression: −0.0055) and based on the downscaled ERA-interim model as provided by Chelsa (*t*_6_ = −24.03, *p* < 0.001, *r*^*2*^ = 0.99, slope of regression: −0.0058, Fig. 2b) (Karger, Conrad, Böhner, Kawohl, Kreft, Soria-Auza, & Kessler, 2017; Karger, Conrad, Böhner, Kawohl, Kreft, Soria-Auza, Zimmermann, et al., 2017). The mean of the mean seasonal soil temperatures (10.88 °C) measured on the 30 plots at our microclimatic heterogeneous pasture was similar to the temperature as predicted by the regression model (Fig. 2b). However, variation between the plots was pronounced (95% confidence interval: 9.67 – 12.33). This range of temperature corresponds to temperature differences expected in elevations differing in 484 or 474 m following the regression models based on our temperature logger or Chelsa data, respectively (Fig. 2b).

### Flowering plants, insects and their interactions

The total plant cover within the plots ranged between 70 % and 100 % (mean ± SD = 90.93 ±7.24). A total of *n* = 59 entomophilous forb species were in anthesis during the observation period. The number of flowering entomophilous forbs (species richness) varied from *n* = 10 to *n* = 23 species per plot (mean ± SD = 16.5 ± 3.41). The number of insect families per plot interacting with flowers (insect family richness) varied from *n* = 3 to *n* = 20 (mean ± SD = 8.77 ± 3.84). The sum of insect interactions with flowers per plot was between *n* = 5 and *n* = 80 (mean ± SD = 32.60 ± 18.22). Interactions between flowers and their visitors were on average rather generalized (network specialization index *H*_*2*_*’*: mean ± SD = 0.37 ± 0.20).

### Correlations between microclimatic heterogeneity, species composition and richness, and flower-visitor interactions

Plant species composition varied between plots and responded to variation in mean seasonal soil temperature, which means that plots with a similar mean seasonal soil temperature were also similar in plant species composition (Mantel test: *r* = 0.21, *p* < 0.01, Fig. 3).

**Fig. 3.**
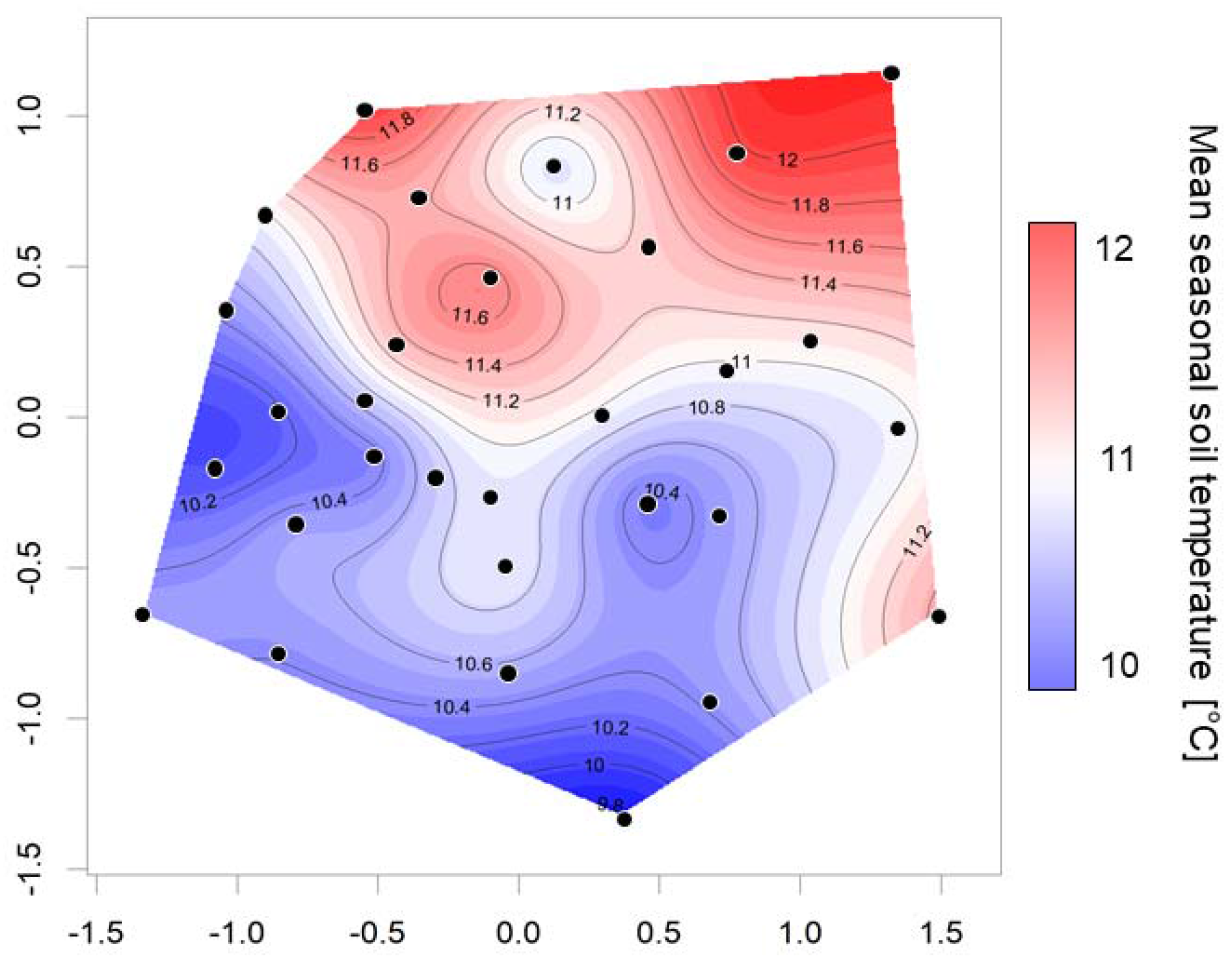
Similarity of study plots in flowering plant species composition in relation to seasonal mean soil temperature. Non-metric multi-dimensional scaling (NMDS) of the dissimilarity (Bray-Curtis) in plant community composition (i.e. beta-diversity) between *n* = 30 investigated 1.5 × 1.5 m plots on an alpine pasture located at 2,273 m a.s.l. in the mountain range of the Hohe Tauern in the Austrian Alps, showing the relationship with mean soil temperature (Mantel test *r* = 0.21, *p* < 0.01). The points resemble the sampled plant communities. The distance between the plots is a measure for the dissimilarity in the community composition (i.e. the further two points are apart the more they differ). Red colors indicate warmer mean seasonal soil temperatures and blue colors resemble colder temperatures.

Mean seasonal soil temperature directly and positively affected the number of flowering plant species of plots (Pearson’s product-moment correlation: *t*_*28*_ = 2.44, *p* = 0.021) as well as plant cover (*t*_*28*_ = 2.83, *p* < 0.01, Fig. 4). The number of insect families increased with increasing plant species richness (*t*_*28*_ = 2.98, *p* < 0.01), but not directly with mean seasonal soil temperature (*t*_*28*_ = 0.22, *p* = 0.828). The total number of flower-insect interactions was positively correlated with both higher plant species richness (*t*_*28*_ = 2.28, *p* = 0.030) and insect family richness (*t*_*28*_ = 11.96, *p* < 0.001). Note, that the number of interactions per plant or animal taxon was unaffected by the number of plant or animal taxa present on plots (*t*_39_ ≤ 1.75, *p* ≥ 0.088), thus interactions per taxon remained on average the same independent on community context. Complementary specialization of networks *H*_*2*_*’* was unaffected by both plant (*t*_*27*_ = −0.82, *p* = 0.420) and insect richness (*t*_*27*_ = −0.67, *p* = 0.507). However, we found a higher generalization (i.e. lower *H*_*2*_*’*) of flower-visitor networks on plots with warmer mean seasonal soil temperatures and a higher plant cover (all results are summarized in Fig. 4). Apart from the correlations described above, no other correlations between the parameters shown in Fig. 4 were significant (Supplementary Table 1). Correlation network analysis based on Pearson’s coefficient of correlation *r* and considering significant correlations only, revealed that mean seasonal soil temperature and plant species richness had the highest degree (*DC*) and betweenness centrality (*BC*) (Fig. 4). These results are largely supported by a structural equation model (Supplementary Information 1).

**Fig. 4.**
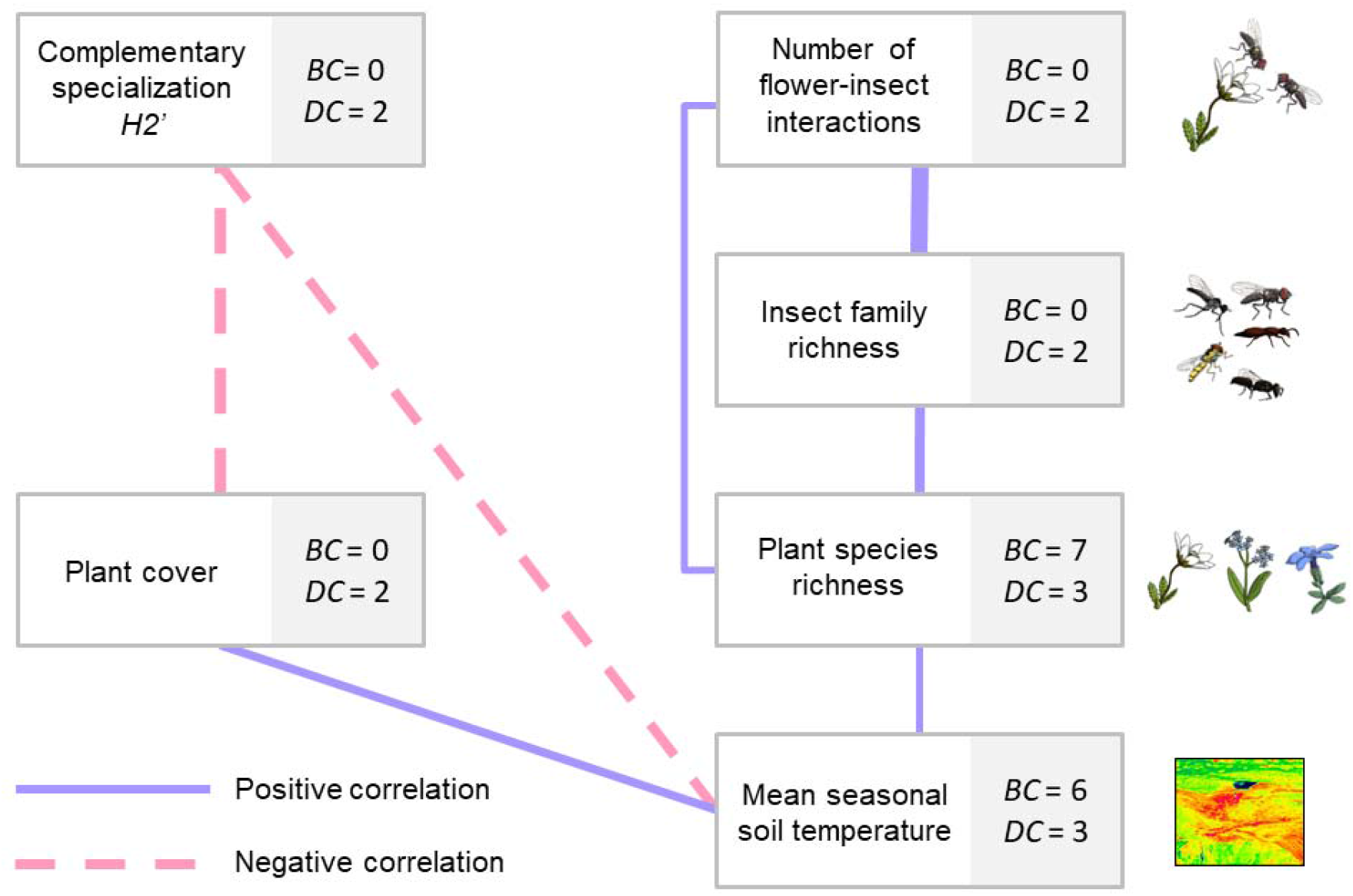
Relationship between mean seasonal soil temperature and parameters characterizing plant and animal communities as well as their interactions. Correlation network based on Pearson’s product-moment correlations showing significant relationships between mean seasonal soil temperature of *n* = 30 plots (2.25 m^2^) on an alpine pasture, plant cover, species richness of flowering plants, number of insect families recorded on flowers, number of observed insect interactions with flowers and complementary specialization *H*_*2*_*’* of flower-visitor interaction networks. Blue lines indicate positive correlations, red dashed lines indicate negative correlations. Edge width is proportional to Pearson’s r. Betweenness (*BC*) and degree centrality (*DC*) are shown for each parameter.

## Discussion

Microclimatic heterogeneity is particularly pronounced in alpine environments (Scherrer & Körner, 2010) and has been suggested to affect the diversity and composition of plant communities (Opedal et al., 2015). However, whether microclimatic effects also affect higher trophic levels, such as flower visitors, remains unknown. In our study area we detected strong microclimatic heterogeneity with differences in soil temperature of 2.8 °C between our study plots. Plant communities responded to increased temperatures with higher cover and higher species richness in flowering plants. Next to species numbers, microclimate also affected plant species composition with different species assemblages on plots with different mean seasonal soil temperatures. These effects of microclimate on the alpha- and beta-diversity of plant communities lead to a higher richness of insects consuming floral resources on warmer plots. No direct effect of soil temperature on insect visitors was evident. Therefore, diversity of higher trophic levels is suggested to be indirectly bottom-up controlled by microclimate. Additional to the direct microclimatic effects on plant communities and the indirect effects on insects, microclimatic heterogeneity also affects the structure of interaction networks with more generalized interactions on warmer plots than on colder ones.

The strong microclimatic heterogeneity of mean seasonal soil temperature is mostly attributed to daily temperature maxima. The magnitude of temperature variability and therefore heterogeneity is especially pronounced during midday, when the highest soil temperatures are reached. Therefore, aspect, slope and shading of the plots resulting in differences in the intake of solar irradiance were the main cause of microclimatic heterogeneity. Further factors such as air temperature, wind speed, surface albedo, microtopography, substrate type, and evapotranspiration may additionally increase temperature variability (Graham et al., 2012; Scherrer & Körner, 2010; Wundram et al., 2010). In concordance to earlier findings, we found a higher vegetation cover on warmer plots compared to colder plots, which may be attributed to a positive relationship between plant productivity and temperature demonstrated in several studies (Anderson & McNaughton, 1973; Kikvidze et al., 2005; Lembrechts et al., 2018). Additionally to plant cover, higher temperatures also positively affected species richness of insect pollinated plants (see also (Schöb, Kammer, Choler, & Veit, 2009)) and also affected plant community composition. Thus, microclimatic heterogeneity may offer more temperature niches that environmentally filter different assemblages of plant species (Bagousse□Pinguet et al., 2017; Bello et al., 2013; Blonder et al., 2015; Kraft et al., 2015; Myers & Harms, 2011), which leads to a larger number of plants that are able to establish in a given area (Conti et al., 2017; Löffler & Pape, 2008; Lundholm, 2009; Opedal et al., 2015). Thus, alpha-diversity of plants in an alpine habitat is facilitated by a high small-scale beta-diversity, i.e. by changing species assemblages due to spatial variations in soil temperature.

Additional to the finding that microclimatic heterogeneity affects plant species assemblages, we demonstrated that also higher trophic levels (e.g. flower visiting insects) and interactions between trophic levels respond to microclimate. Enthomophilous plants depend on insects for reproductive success and, in turn, flower visiting insects depend on flowering plants for resources suggesting a tight dependency between these trophic levels, potentially even on smaller scales. Flower visiting insects are believed not to be directly influenced by soil temperature because the activity of flower visitors is mainly affected by larger scale weather conditions, such as air temperature, solar radiation and wind (Kühsel & Blüthgen, 2015; Totland, 1994). Correspondingly, we found no direct effect of local soil temperature on the richness of flower visitors. However, we found an indirect bottom-up effect of microclimate on the number of flower visiting insect families, which increased with higher flowering plant species richness found at warmer plots. Species richness of insect communities has been shown to be closely related to floral diversity (i.e. resource diversity, (Ebeling et al., 2008; Potts, Vulliamy, Dafni, Ne’eman, & Willmer, 2003). This finding may be explained by the fact that different species of flower visitors exhibit a specialization to specific floral traits (Junker et al., 2013). Communities with a larger number of plant species are assumed to offer more niches that can be occupied by a higher number of flower visiting insect species specialized to specific flower traits (Junker et al., 2015). Furthermore, higher plant species richness and a higher number of insect families led to more insect-flower interactions in warmer plots. This finding results from an additive effect of more interacting taxa instead of more frequent interactions per taxon because the number of interactions per taxon was unaffected by taxon richness per plot, which may explain the positive correlation between plant and animal taxa. Interestingly, not only richness of plant and animals are directly or indirectly were affected by microclimate, also the interaction structure between these trophic levels varied with mean seasonal soil temperature. Interaction networks were more generalized (i.e. lower *H*_*2*_*’*) on warmer plots that also featured a higher plant cover. While we have no clear mechanistic explanation for this finding, these results are however suggesting that warmer plots favour productivity (higher plant cover) (Kikvidze et al., 2005; Lembrechts et al., 2018) and the stability of the plant and pollinator communities (lower *H*_*2*_*’*) (Blüthgen & Klein, 2011). The relationships between plant diversity and productivity (Fraser et al., 2015) as well as between plant diversity and community stability (Blüthgen & Klein, 2011) are well established. In contrast, the relationship between productivity and community stability has been suggested, only (Worm & Duffy, 2003). Our results therefore support the notion that more productive systems may also be more stable – mediated by higher mean seasonal soil temperature – even on a small scale, and therefore may stimulate more research in this direction in the future. Overall, correlation network analysis assigned high degree and betweenness centrality to mean seasonal soil temperature and plant species richness suggesting a central role of these parameters in mediating the productivity, species richness across trophic levels, and interaction structure and stability of communities.

Plant–insect interactions are increasingly affected by climate warming possibly leading to temporal and spatial mismatches among mutualistic partners (Hegland et al., 2009). Temperature is the main trigger of flowering phenology for alpine species and local variations in temperature can lead to a shift in flowering phenology (Cleland, Chuine, Menzel, Mooney, & Schwartz, 2007; Hülber, Winkler, & Grabherr, 2010), which may be particularly relevant in alpine landscapes (Carbognani, Bernareggi, Perucco, Tomaselli, & Petraglia, 2016; Inouye, 2008; Jackson, 1966). Next to phenology shifts and associated desynchronization between plants and their flower visitors, upward migration of plants to find suitable temperature niches (Alexander, Diez, & Levine, 2015; Grabherr, Gottfried, & Pauli, 1994; Pauli et al., 2012) may disrupt plant-visitor interactions. The observed mean seasonal temperature difference of 2.8 °C between our microclimatic sites that were located in close vicinity and on the same elevation corresponds to the temperature difference as measured between sites that differ in nearly 500 meters in elevation, which is also in agreement with an average decrease in air temperature of 0.6 C° per 100 meters upslope (Körner, 2007). The extent of temperature variation provided by microhabitats due to local small-scale topography found in this study corresponds to the most likely warming scenarios for the next 100 years of 3 °C as reported in the Intergovernmental Panel on Climate Change (IPCC) projections. Therefore, the pronounced microclimatic heterogeneity suggests that microclimatic habitats are capable of providing sufficient heterogeneity to buffer climate change impact on alpine plants (Graham et al., 2012; Lenoir et al., 2013; Scherrer & Körner, 2011) and on higher trophic levels dependent on plant resources. Thus, microclimatic heterogeneity may reduce the necessity of upward migrations of species as a response to climate warming. In conclusion, alpine plants and their consumers can experience the same climatic conditions in climate refuges within a few meters distance on the same elevational level. Therefore, migration of a few meters may be sufficient to track their suitable climate niche.

Our study provides evidence that heterogeneity in mean seasonal soil temperature has the potential to promote flowering species richness of alpine plants (alpha-diversity) at local scales by facilitating the beta-diversity among different microclimates. Our results show that the positive effect of soil temperature on plant cover and richness of flowering plants is carried on to the next trophic levels by increasing the richness in these taxa and also by affecting the network structure between trophic levels. Microclimatic heterogeneity may thus have the capability of mitigating climate warming impacts on alpine plants and their interactions with flower visitors by buffering possible mismatches in phenologies *via* desynchronized flowering at different temperatures and by increasing overall diversity, leading to more stable systems.

## Acknowledgements

We thank the Haus der Natur and the National Park Hohe Tauern for logistic support, and Andreas LaFolette for help in the field and Florian Griessenberger for help in statistical analysis. The study was funded by the Austrian Science Fund (FWF, P29142-B29).

## Author Contributions

RRJ conceived the study. RRJ, LMO, ML designed the study. LMO and ML conducted fieldwork. LMO, RRJ performed statistical analyses. LMO and RRJ drafted the manuscript and all authors contributed to the final version.

## Data accessibility

The data supporting the results can be found in Supplementary Table 2–5.

## Supplementary Information

### Supplementary Information 1

#### Effect of microclimatic heterogeneity on plant-pollinator interactions

Additionally to the analysis presented in the main text we tested the effect of microclimatic heterogeneity on plant-pollinator interactions using a structural equation model (Supplementary Information 1 Fig. 1). Based on our main hypothesis that the seasonal mean temperature affects plant and animal communities as well as their interactions and based on pre-existing knowledge from the literature and own experience, we hypothesized causal relationships between seasonal mean seasonal soil temperature of the plots, plant cover, the species richness of flowering plants, the number of insect families recorded on flowers, the number of observed insect interactions with flowers and complementary specialization *H*_*2*_*’* (Blüthgen, Menzel, & Blüthgen, 2006) of flower-visitor interaction networks. We hypothesized that our main effect, the seasonal mean temperature of the plots, directly affects the four other variables. We further hypothesized direct effects of plant richness on insect family richness and complementary specialization *H*_*2*_*’;* and of insect family richness on complementary specialization *H*_*2*_*’*. We fitted the structural equation model based on these hypotheses to the field data using default settings of the R package *lavaan* and a maximum likelihood (ML) estimator (Rosseel, 2012).

Mean seasonal soil temperature directly and positively affected number of flowering plant species of plots. Whereas mean seasonal soil temperature was not directly related to the number of insect families that visited the flowers within a plot, an indirect effect is indicated by the positive effect of number of flowering plant species on number of insect families. Flower abundance varied independently on mean seasonal soil temperature and number of flowering plant species. Complementary specialisation of networks *H*_*2*_*’* was unaffected by plant and insect richness as well as by flower abundance. However, we detected a negative relation between mean seasonal soil temperature and *H*_*2*_*’*.

**Supplementary Information 1 Fig. 1.**
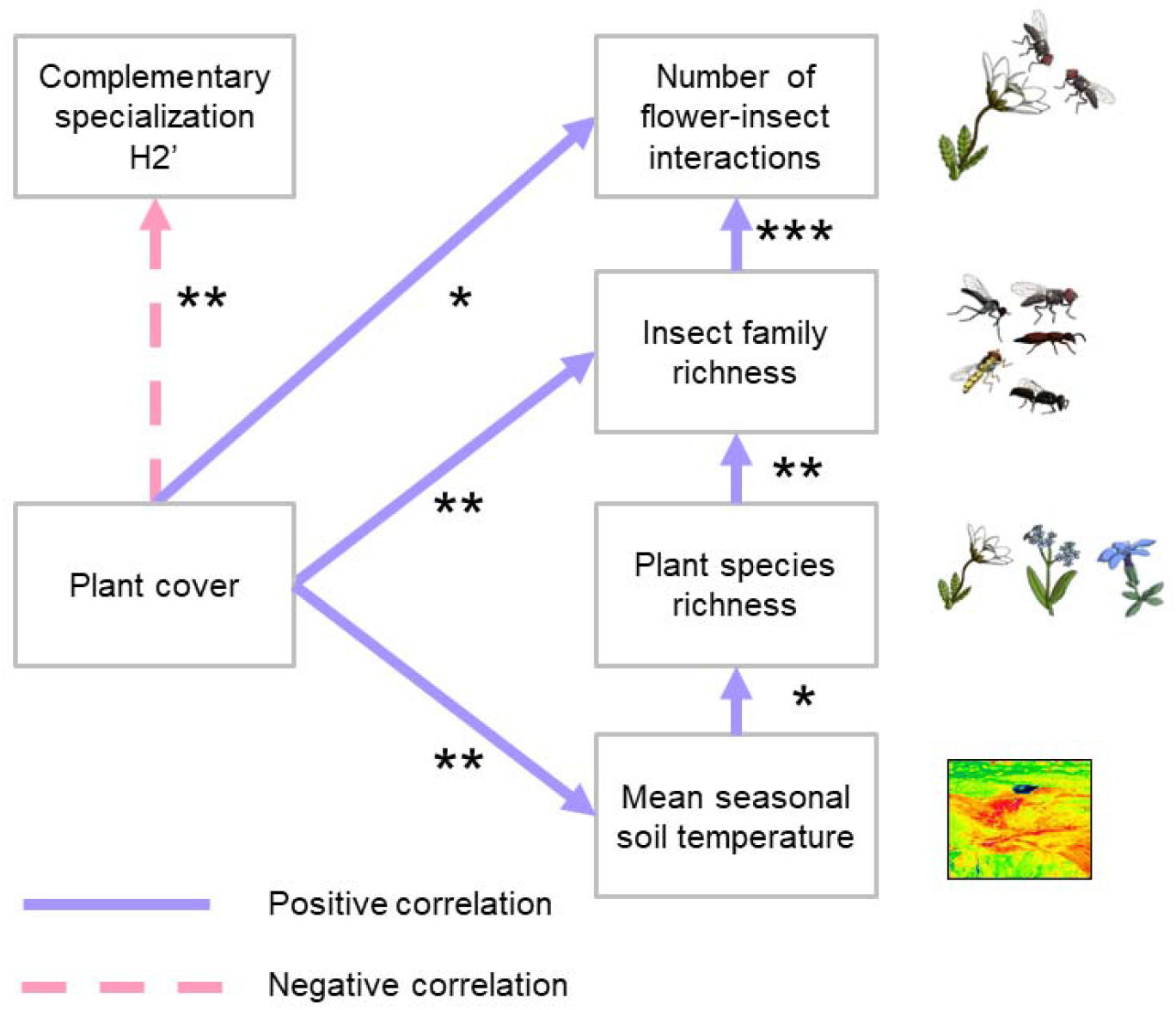
Structural equation model of mean seasonal soil temperature and parameters characterizing plant and animal communities as well as their interactions. Structural equation model showing the relationships between mean seasonal soil temperature, plant cover, species richness of flowering plants, number of insect families recorded on flowers, number of observed insect interactions with flowers and complementary specialization *H*_*2*_*’* recorded on *n* = 30 plots (2.25 m^2^) on a topographically heterogenous alpine pasture located at 2,273 m a.s.l. in the mountain range of the Hohe Tauern in the Austrian Alps (12°49’21” E 47°07’24” N). Arrows indicate significant correlations between the tested parameters (blue: positive correlation, pink: negative correlation).

